# Ecological network analysis reveals cancer-dependent chaperone-client interaction structure and robustness in the mitochondria

**DOI:** 10.1101/2022.11.24.517852

**Authors:** Geut Galai, Barak Rotblat, Shai Pilosof

## Abstract

Cancer cells alter the expression levels of metabolic enzymes to fuel proliferation. The mitochondrion is a central hub of metabolic reprogramming, where multiple chaperons service hundreds of clients, forming complex chaperone-client interaction networks (CCINs). Studying (i) how network structure varies across cancer types and (ii) the influence of structure on networks’ robustness to chaperon targeting is key to developing cancer-specific drug therapy. We investigate these yet unaddressed questions using theory and methodology from network ecology. We show that CCINs are non-randomly structured across cancer types. Moreover, we identify groups of chaperones that interact with similar clients, providing specialization on clients but also redundancy in the case of chaperone removal on the other. Simulations of network robustness show that this modular structure strongly affects cancer-specific response to chaperon removal. Our results open the door for new hypotheses regarding the ecology and evolution of CCINs and can inform caner-specific drug development strategies.

## Introduction

Cancer cells reprogram their metabolism to fuel proliferation, promote undifferentiated state, drive metastasis, overcome metabolic stress and communicate with the immune system [1]. Metabolic reprogramming is a hallmark of cancer and is executed by changes in the expression levels of metabolic enzymes localized to the cytosol or the mitochondria. The mitochondria is a major metabolic hub playing a crucial role in metabolic reprogramming. The vast majority of mitochondrial proteins are synthesized in the cytosol and imported into the mitochondria in an unfolded state after which they are folded by mitochondrial chaperones [2]. There are fifteen mitochondrial chaperones, including ATP-dependent proteases, catering to the folding and elimination of damaged and misfolded mitochondrial proteins [3]. In cases where demand for chaperone activity is higher than the available folding capacity, proteins will misfold, aggregate and be degraded [2]. Weather specific proteins are clients of specific chaperones and dependent upon them for their folding and degradation in the mitochondria is not known. Since metabolic reprogramming entails upregulation of particular mitochondrial enzymes, it is important to know if targeting a specific mitochondrial chaperone will lead to the collapse of its particular protein substrates.

The dependency of tumor cells on particular mitochondrial chaperones is exemplified by the biological activity of small molecules targeting mitochondrial chaperones such as TRAP1 [4], HSPD1 [5], ClpX [6] and ClpP [7], all of which are promising anti-cancer compounds. Interestingly, it is not clear why particular cell types are sensitive to a chaperone’s particular inhibitors. One possibility is that the dependency of a client on a chaperone is dictated by environmental factors such as different cancer types. In that case, targeting a specific chaperone will lead to the collapse of different clients in different cancer types and tumor entities, resulting in cancer-specific cellular fates. Another possibility is that different cancer types can tolerate the misfolding of different mitochondrial proteins. In this case it is expected that inhibition of a specific chaperone will lead to the collapse of similar proteins in different tumor entities but the outcome will be specific to a cancer type [8]. Alternatively, if chaperones interact with different clients in different cancers, chaperone inhibition will result in cancer-specific protein misfolding. Addressing these hypotheses requires a framework that can handle the complexity of multiple chaperone-client interactions and how the structure of these networks vary across cancer types.

Ecological network analysis is particularly suitable for analyzing variation in interactions across environments [9]. In parallel, it has been argued that ecological theory can benefit cancer studies [10–12]. In this regard, the study of chaperone-client interactions (CCIs) is largely analogous to the study of ecological species interactions networks. chaperone-client interactions, like, for example plant-pollinator interactions, form bipartite networks which describe links between two distinct sets of nodes [13]. In ecology, the environment is a strong determinant of species interactions [9,14]. Therefore, even when two species co-occur in multiple environments, they may interact in one environment and not the other [15,16]. Analogously, cancer types are distinct environments that may modulate CCIs. The null hypothesis is that if the interaction between a chaperone and its client is enabled only by their biophysical traits alone, they should interact in every cancer environment. The alternative hypothesis is that variation among cancers expressed in different tissues implies variation in metabolic demands and reprogramming, which should result in a concomitant variation in CCIs, and therefore network structure, across cancers.

Testing this hypothesis can provide insights into how variation in interactions translates to variation in system stability. The structure of ecological networks largely affects their stability and function [17]. For instance, in mutualistic networks, redundancy in interactions, whereby the same plant is pollinated by multiple insects increases network resilience because when one pollinator goes extinct, others can still pollinate the plant [18]. Modularity—a structural pattern in which the network is partitioned into groups of species that interact densely among themselves and sparsely with others [19]—increases network stability as it slows down perturbations by confining them to modules [20]. Similar phenomena can occur in CCI networks, but this has not been tested. Delineating the interplay between structure and stability in different cancer environments in CCIs is crucial because controlling metabolic reprogramming via inhibitors that target specific chaperones may be cancer-specific.

Here, we apply ecological network analysis to a set of CCI networks from 12 cancer environments. Given the interdisciplinarioty of this manuscript we provde a gloassary of terms (Table 1) The chaperones and their clients occur in all the networks, allowing us to investigate the effect of cancer environment on interaction redundancy and structure. We specifically ask: (1) How do CCIs vary across cancers and what might cause this change in cancers? (2) What are the consequences of this variation to the stability of the network in each cancer environment following removal of chaperones. We reveal a non-random and hierarchical pattern by which cancers modulate the chaperones’ ability to realize their client interaction potential. Moreover, there is strong niche separation whereby groups of chaperones interact with distinct clients. Nevertheless, there is still redundancy such that chaperones interact with similar sets of clients, increasing the robustness of the networks to targeted removal of chaperones. Niche separation, redundancy and robustness vary across cancer environments, highlighting the role of cancer type in modulating chaperone-client interactions.

**Table 1:** Glossary. Definitions of key terms used in this paper.

| Term | Definition |
| --- | --- |
| Chaperon | A mitochondrial protein that is part of the agents supporting proteostasis in the mitochondria [3]. |
| Client | A mitochondrial protein with a potential interaction with a chaperone. |
| Ecological network | A mathematical graph object that represents pairwise biotic interactions (edges)—such as competition, predation, pollination—between multiple species (nodes) [21]. Networks can be depicted as matrices in which each row and column represent an entity and matrix entries are the links. |
| Niche | The set of biotic and abiotic conditions within which an organism can perform biological functions. In the context of ecological interactions, for example, hosts are the niches of parasites. Here, we consider a chaperone’s clients as its niche. |
| Niche overlap | The degree to which two species overlap in the set of conditions they can exploit. For instance, the common set of prey of two predators. Here, the set of clients that interact with two chaperones. |
| Niche partitioning | The processes that drive two species to minimize overlap in their niches (e.g., due to competition). Here, the tendency of two chaperones to interact with different sets of clients. |
| Nestedness | A particular network organization in which the more specialist nodes interact with proper subsets of the more generalist ones [13]. |
| Realized niche | The exploitation of a subset of all potentially exploitable resources due to some environmental or biotic constraints. Here, the realized niche of a chaperon are the subset of the clients with which it interacts. |
| Specialization | The extent to which a species can exploit resources out of all available ones [22]. Here, the number of clients a chaperone interacts with out of all clients. |

## Results

### Chaperone interaction patterns are non-randomly affected by cancer type

Our networks encode interactions between 15 mitochondrial chaperones and 1, 142 client proteins across 12 cancer environments. We estimated interactions based on co-expression data while normalising sample size across cancer types using bootstrapping (Methods; Table S1). All chaperones and clients were present in all 12 networks. We defined *P_c_* as the total number of clients a chaperone *c* can interact with across cancers. We calculated the level of specialization *S_c_* of each chaperone *c* as the total number of clients it interacts with (across all cancers) divided by the 1,142 potential protein clients, *S_c_* = *P_c_*/1,142. A value of 1 indicates that the chaperone can interact with all 1,142 proteins. Specialization reflects variation in chemical and physical properties that enable chaperones to interact with mitochondrial proteins, independently of cancer. In ecological jargon, chaperones with low values of *S_c_* are broadly considered specialists and those with high values as generalists. Specialization ranged from 40%-65% (55.5 ± 8.1%) (Fig. S1A). Because in our system all clients were present in all cancers, the number of clients a chaperone interacts with should be equal across cancers if only biophysical properties determine interactions. Deviation from uniformity for a given chaperone indicates that the cancer type affects chaperone’s interactions. To test this, we calculated cancer-specific specialization—the proportion of proteins a chaperone interacts with in a given cancer environment—as 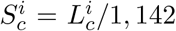, where 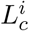 is the number of links of a chaperone *c* in cancer type *i*. Instead of uniform specialization, we find large variability in chaperones’ interactions across cancers (Fig. S1B).

Following the fact that chaperones interact with different sets of clients and considering the evidence for the effect of cancer type on chaperone interactions, we calculated the number of clients a chaperone interacted with in a specific cancer out of all the clients it could potentially interact with, 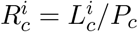. This is analogous to the concept of realized niche in ecology—that is, the exploitation of a subset of all potentially exploitable resources (clients) due to some environmental or biotic constraints. We find a clear pattern in the way cancer types affect the realized niche of chaperones (Fig. 1). From the chaperone perspective: cancer environments affect the realized niche of chaperones. For example, SPG7 interacts with a high proportion of the clients that it can in thyroid cancer (THCA) but a low proportion in breast cancer (BRCA). From the cancer perspective: chaperones vary in their realized niche within the same cancer type. For instance, in BRCA the CLPP chaperone realizes about 40% of its potential, while SPG7 realizes about 15%. Put together, these two observations create a weighted-nested pattern [23,24]. That is, chaperones that interact with a small proportion of their potential clients are subsets of those that interact with a higher proportion (nestedness across rows); on the other hand, cancer environments that enable chaperones to interact with few of their potential substrates are subsets of those that enable higher proportions (nestedness across columns). The pattern of weighted-nestedness was non-random when compared to 1,000 counterpart networks assembled from networks in which chaperone-clients interactions were shuffled [24] (Methods; Fig. S2). Moreover, we found a similar non-random weighted nestedness in the level of specialization across cancers, 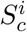 (Fig. S1B).

**Fig. 1:**
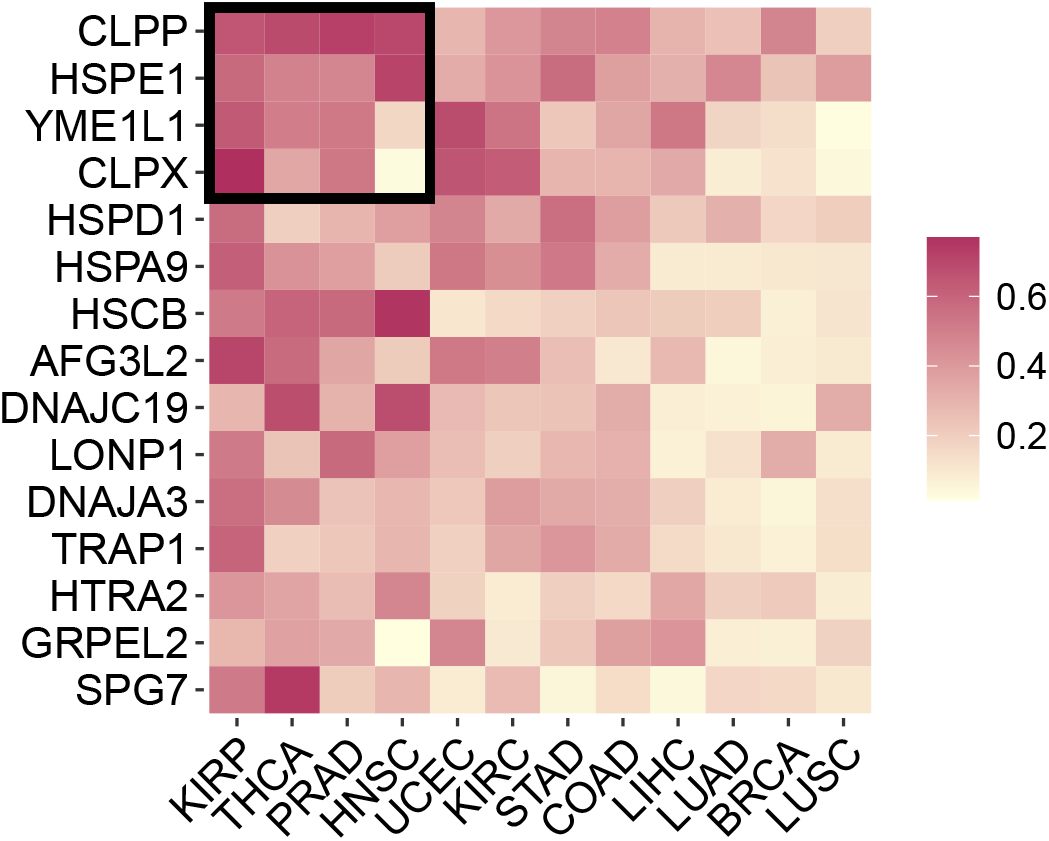
Non-random patterns in chaperone realized niche across cancer environments. Each square depicts the number of clients a chaperone interacts with in a specific cancer out of all the clients it could potentially interact with (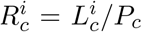; realized niche, see text). Cancer environment affect the realized niche of chaperones as is evident from the non-uniform colors in each row. Chaperones vary in their realized niche within the same cancer type, as is evident from the non-uniform colors in each column. The matrix is significantly weighted-nested, with a core of chaperones with high realized niche values in particular environments. A core of four cancer type and four chaperones (arbitrary selection) is depicted as an example by the black square. Rows and columns are arranged by their marginal sums. A similar weighted-nested pattern was found for chaperone cancer-specific specialization (Fig. S1B)

Taken together, we find non-random structured variation in the way cancer environments mediate chaperone interactions. Specifically, there is a distinct hierarchy in the levels of specialization and realized niche of chaperones on the one hand, and in the way cancers mediate that chaperone interactions on the other. This creates a core of chaperones that interact with most of the substrates that they can in a few cancer types (Fig. 1). We note, that unlike in ecological networks [13,14,24,25], here nestedness is not a pattern of CCIs. Instead, it is a pattern in the realization of interactions across environments.

### Protein and chaperone expression do not explain cancer-mediated interaction patterns

Nestedness could result from variation in protein expression levels across cancers. That is, when the expression of proteins is high, then chaperone expression should also be high to ensure properly supporting of all their clients. This positive correlation in expression should result in more interactions and concomitantly in higher specialization and realized niche values (also see Discussion). To test this hypothesis we compared the distributions of median protein expression values of the different cancers. The distributions were largely similar between cancers (Fig. S3A), implying that hierarchy in realized niche values is not the result of variation in protein expression across cancers. From the chaperones’ perspective, variation in expression should result in variation in their realized niche. Although chaperones did vary in their expression levels across cancers, there were no significant correlations between expression level and realized niche values 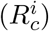 (Fig. 2, S3B, Table S3). Overall, our data indicates that expression levels of chaperones and their clients cannot explain observed patterns in the realized niche and specialization. One alternative hypothesis is that there is a gradient of dependencies on the support for mitochondrial proteins between the different cancers, where some cancers are highly dependent (eg. KIRP) and others less (LUSC).

**Fig. 2:**
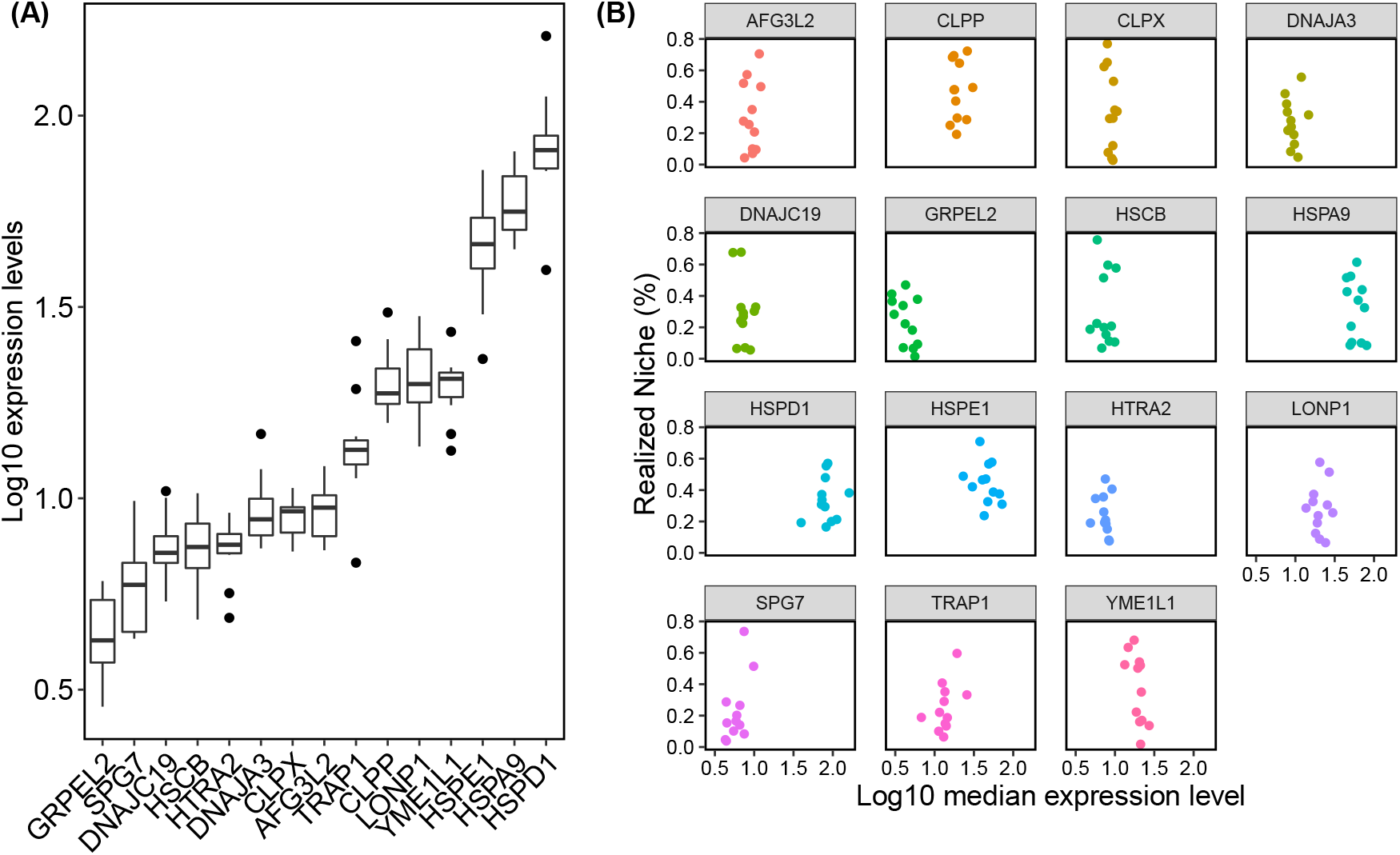
chaperone expression levels. **(A)** The log-transformed median values of chaperone expression levels across cancers. Some chaperones are more expressed than others. **(B)** Each scatter plot presents the distribution of a single chaperone’s realized niche over its log10 gene expression across cancers (each data point is a cancer type). Spearman correlations combined with Bonferroni corrections between these two indices result in non-significant p-values for all chaperones (see Table S3).

### chaperones interact with similar clients in different cancer environments

The strong and non-random variation in specialization and realized interactions (Fig. 1) does not provide the full picture because this analysis ignores the identity of the clients. To further test the effect of cancer type on chaperone interactions, we used the Jaccard similarity index to compare the identity of the clients of each chaperone *c* between pairs of cancer types *i* and 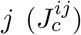. We then calculated the partner-fidelity of each chaperone—an ecological measure of how repetitive are the interaction partners of a species in different places [26]—as the mean of 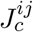 (Fig. S4A). Partner fidelity 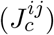 ranges between 0 (a chaperone interacts with completely different clients in each cancer) and 1 (a chaperone interacts with exactly the same clients across cancers). For each chaperone we compared its observed partner fidelity with that expected at random using z-scores (sensu [26]; Methods). We found that although the partner fidelity of chaperones had a median of only 0.25, it was still higher than expected at random for each 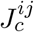 (*z* > 1.96 for all comparisons; also, compare red and blue distributions in Fig. 3A).

**Fig. 3:**
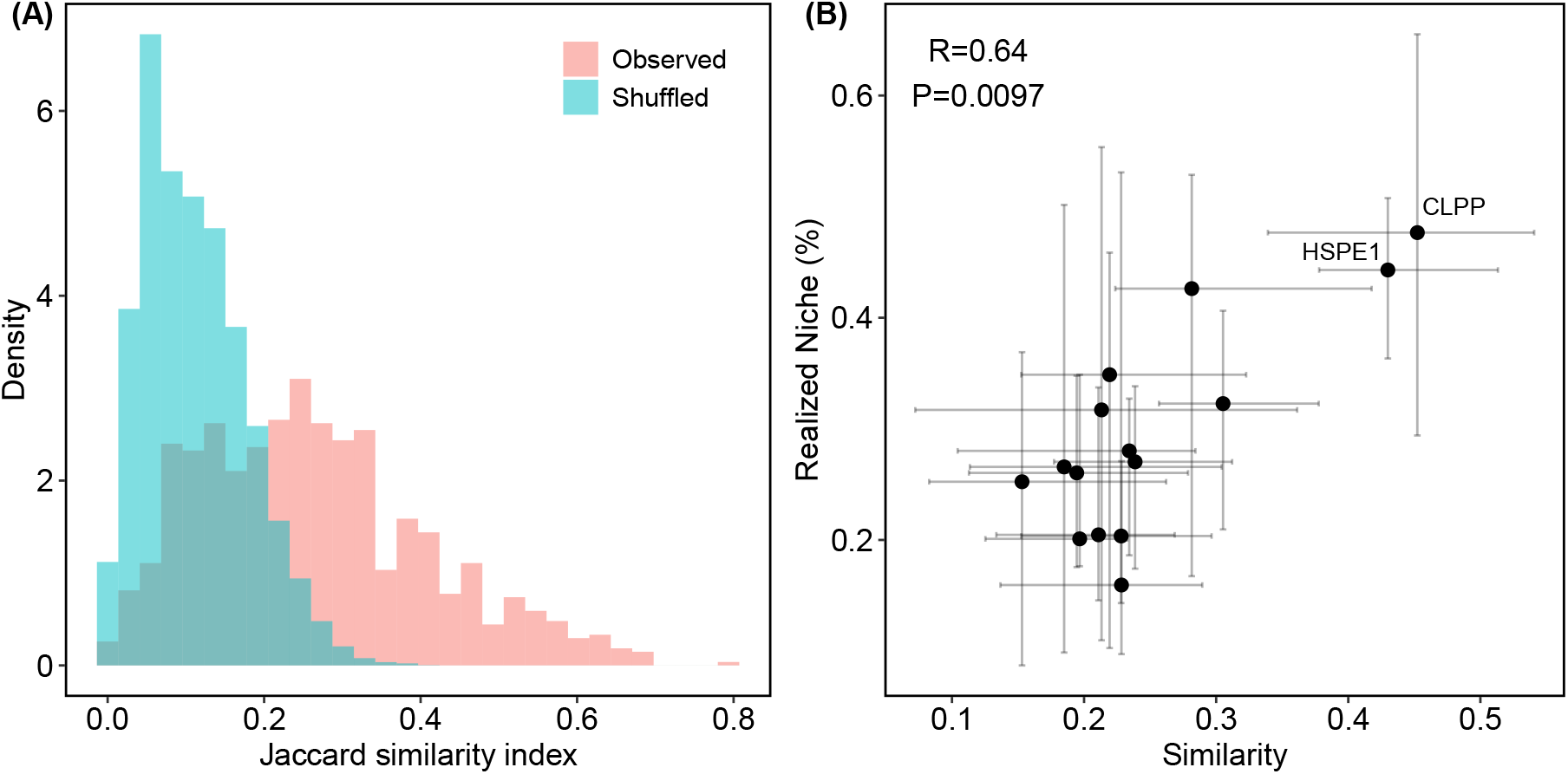
Comparison of client identities between cancers. **(A)** Partner fidelity: the similarity in client identities that a chaperone interacts with between two cancer types calculated as 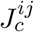 (see text) for observed (red) and counterpart shuffled networks (blue). Overall, chaperones tend to conserve interacting proteins to a limited degree (median 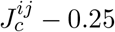) but more than expected at random (median = 0.102). **(B)** These similarity values are positively correlated with the realized niche (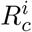. Each data point is a chaperone.

These results indicate that although chaperones tend to conserve their interaction partners across cancers, they can only do so to a limited extent. Hence, client identity could influence specialization and realized niche. To test this, we correlated the median partner fidelity with the median realized niche of each chaperone (Fig. 3B). We find a positive correlation (*r* = 0.64, *P* = 0.0097), which is expected for chaperones with high realized niche values (e.g., CLPP or HSPE1) because interacting with a large proportion of the possible clients inevitably leads to high chances of interacting with the same clients. However, the trend we uncover is not trivial for chaperones with low realized niche values because the few clients being interacted with of the potential clients could be the same or not. The positive correlation indicates that chaperones that tend to interact with few of their potential clients do so with distinct clients across cancers.

### chaperones demonstrate niche separation and redundancy in client interactions

After testing the effect of cancer on chaperone-client interactions, we now turn to explore within-cancer effects on chaperone interactions. We compared the sets of clients that two chaperones *x* and *y* interact with within each cancer *i* using the Jaccard similarity index, 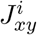 (Methods). 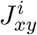 ranges from 0 (the chaperones interact with complete discordant client sets) to 1 (the chaperones interact with exactly the same clients in the same cancer). There was a high occurrence of very low 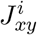 values, which were also lower than expected when compared to shuffled networks (Fig. 4A; See Fig. S4B for within-cancer distributions). This pattern is indicative of ecological niche separation, whereby chaperones tend to interact with different clients. Supporting this claim is the fact that five out of the six chaperones in module 2 are co-chaperones.

**Fig. 4:**
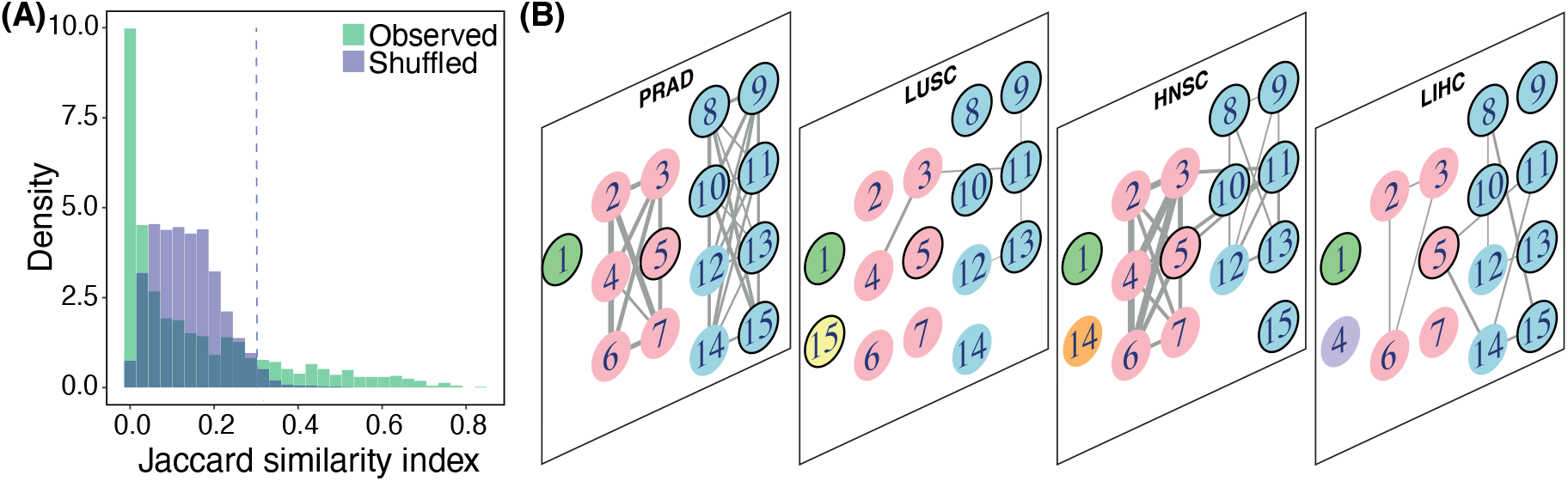
Niche separation in chaperone interaction partners. **(A)** The Jaccard similarity index for each pair of chaperones in a specific cancer (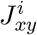; see text). The green and blue histograms depict similarity values across all cancers for the observed and shuffled networks, respectively. Each distribution contains 1260 pairwise comparisons (105 pairwise comparisons in 12 cancer types). There is a mode of lower than random similarity, indicating that in general chaperones interact with sets of clients that are more different than expected at random. However, there is also a tail with chaperone pairs that highly overlap in the clients they interact with. **(B)** A multilayer network of chaperone niche overlap. Each layer is a cancer type. Nodes are chaperones. Numbers insider a node correspond to chaperon ID in Table S2. Nodes with bold and light borders are chaperons and co-chaperones, respectively. Links encode the similarity in clients two chaperons interact with in a cancer 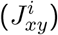—the same values as in panel A. For clarity, only links with weights 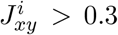 are shown (depicted with vertical line in panel A). Chaperones are divided into six modules (depicted by node color). Except of three instances, each chaperone was clustered into a single module across all 12 cancer types, indicating strong niche separation. For clarity, only four out of 12 cancer types are shown because the division of the chaperones into modules in the missing cancer layers is identical to that of the PRAD cancer.

Niche separation could weaken the robustness of the network to chaperone failure because if a chaperone is not functional then its clients remain unsupported. On the other hand, robustness could be increased if some redundancy exists in client identities, such that when a chaperone is not expressed, another can take its place. Indeed, the distribution of 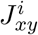 also had a long tail of high values (Fig. 4A; Fig. S4B), indicating that some chaperone pairs interact with highly overlapping sets of clients. Therefore, while the vast majority of chaperone pairs interact with distinct sets of clients, there are still a few pairs with high redundancy, that may provide stability to chaperone removal.

We investigated the interplay between niche separation and robustness by looking for clusters of chaperones and clients that interact more among themselves than with others across cancers using modularity analysis [19,27,28]. In a modular network, chaperones within the same cluster (module) interact with the same clients (redundancy), and modules indicate niche separation. We tested for modularity on all the 12 CCI networks simultaneously, using an ecological multilayer modularity analysis (Methods). In the multilayer network, each layer is a cancer-specific CCI network. Modularity is calculated on all cancers simultaneously [29]. This approach is more adequate than analyzing cancer types separately because it considers all the clients that a chaperone can interact with across cancers, allowing us to detect niche separation and redundancy across all chaperones and cancer types simultaneously. Therefore, we can explicitly include the effect of cancer type on the division to modules. If a module encompasses multiple layers, the cancer type has little effect on the assignment of chaperones to modules. In contrast, if chaperones are assigned to different modules in different cancer types, the effect of cancer is strong.

The multilayer network was partitioned into six modules, of which the chaperones were divided into three main modules across cancers (Figure 4B). chaperones within each module tend to interact strongly with the same clients and weakly with clients from the other modules. A modular structure reflects niche separation between chaperones from different modules on the one hand, but redundancy on the other because chaperones from the same module interact with the same clients. This conclusion is supported by the fact that the chaperones in module 1 are all chaperones and those in module 2 are co-chaperones.

### Network structure affects the robustness of CCIs to chaperone removal

A major goal of studying network structure is to understand its effect on the robustness of the networks to perturbation [17]. A relevant question in our case is how chaperone removal affects client interactions, an analysis that can provide insights into cancer therapy. To explore this questions we conducted an in-silico robustness analysis. While this analysis is common in ecology [18,30], it has not been done in the context of chaperone-client mitochondrial networks. Our algorithm simulates the targeting of particular chaperons and works as follows: in each cancer type, we removed chaperones sequentially. When a client was left without any connected chaperones it was also removed and recorded as a co-extinction. We then plotted the proportion of clients remaining as a function of the proportion of chaperones removed. We calculated a robustness score, *T*, as the area under the extinction graph (Fig. 5) [31]. To explicitly link robustness to network structure we removed chaperones in four scenarios of removal order: (i) from the most to the least connected; (ii) most to least connected within module 1 and then similarly within module 2; (iii) most to least connected within module 2 and then similarly within module 1; (iv) randomly, to obtain a benchmark control comparison within each cancer [18]. Scenarios (ii) and (iii) directly link the modular structure to robustness. We expected that within each cancer, robustness scores in scenarios (i)-(iii) will be lower than the random scenario, and that the network will collapse most rapidly (lowest *T*) in scenario (i).

**Fig. 5:**
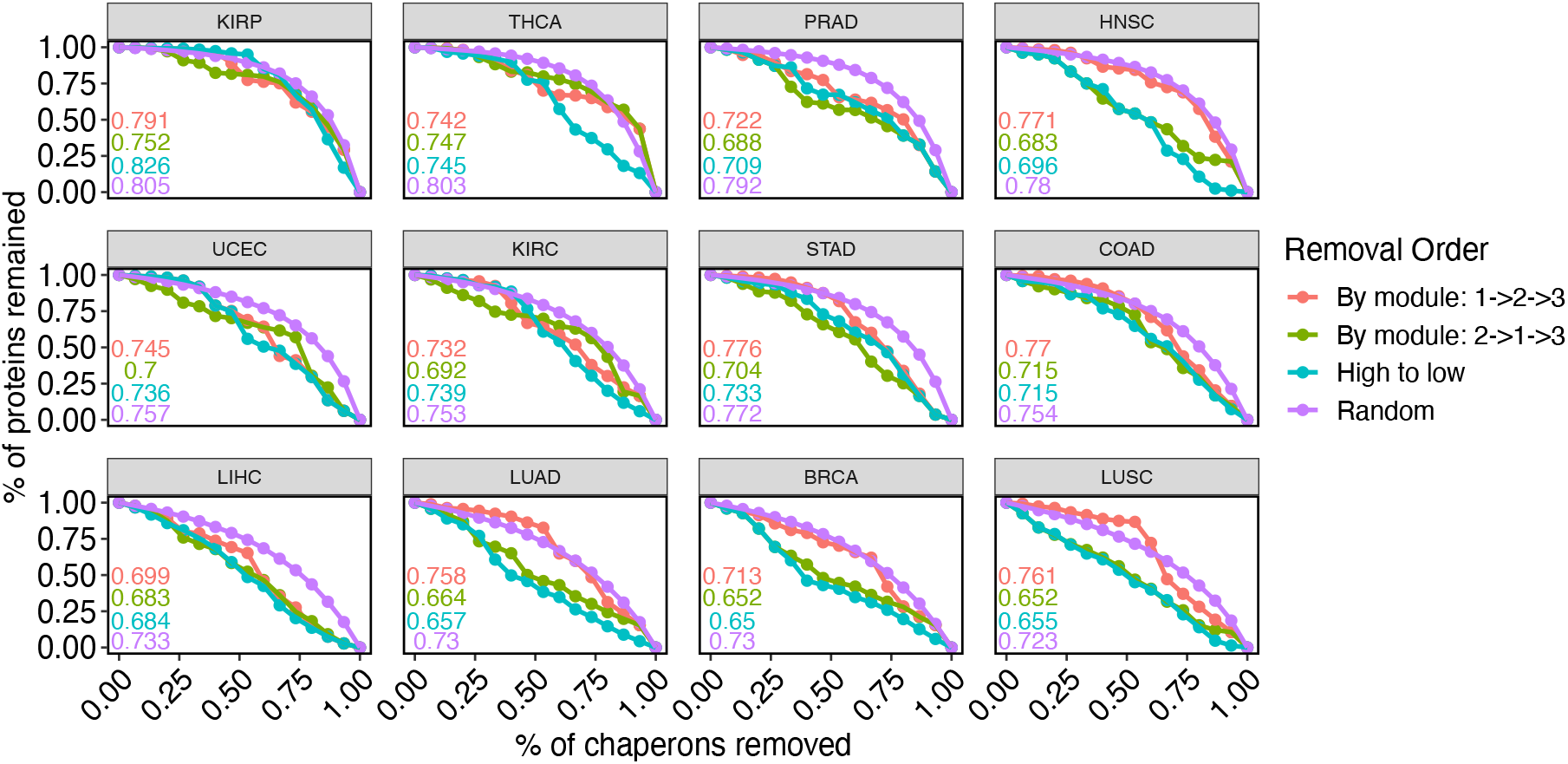
Robustness to chaperone removal. The collapse process of each network can be described by calculating the proportion of clients that remain connected to at least one chaperone, as a function of chaperone removal. There were four scenarios of chaperone removal order. Network robustness *T* is calculated as the area under the curve. Values of *T* are depicted for each scenario in its corresponding color.

In general, the networks collapsed faster in scenarios (i)-(iii) compared to random removal (Fig. 5). However, cancer type strongly affected network collapse. While removal order had little effect on robustness in some cancer types (e.g., COAD, KIRC), in others the effect was strong (e.g., BRCA, LUAD) (Fig. 5). Robustness is tightly linked to the general level of connectivity of the network. Cancer types that increase chaperones’ realized niche value should be more robust (e.g., KIRP) than those in which chaperones realize few of their potential interactions (e.g., LUSC) (Fig. 1). We tested this hypothesis by correlating the mean realized niche values 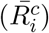 with robustness *T* for each removal scenario. These correlations were statistically significant for all scenarios (Fig. 6). Therefore, the robustness of cancer networks depends both on the general level of connectivity of chaperones and on the mesoscale modular structure of chaperone-client interactions.

**Fig. 6:**
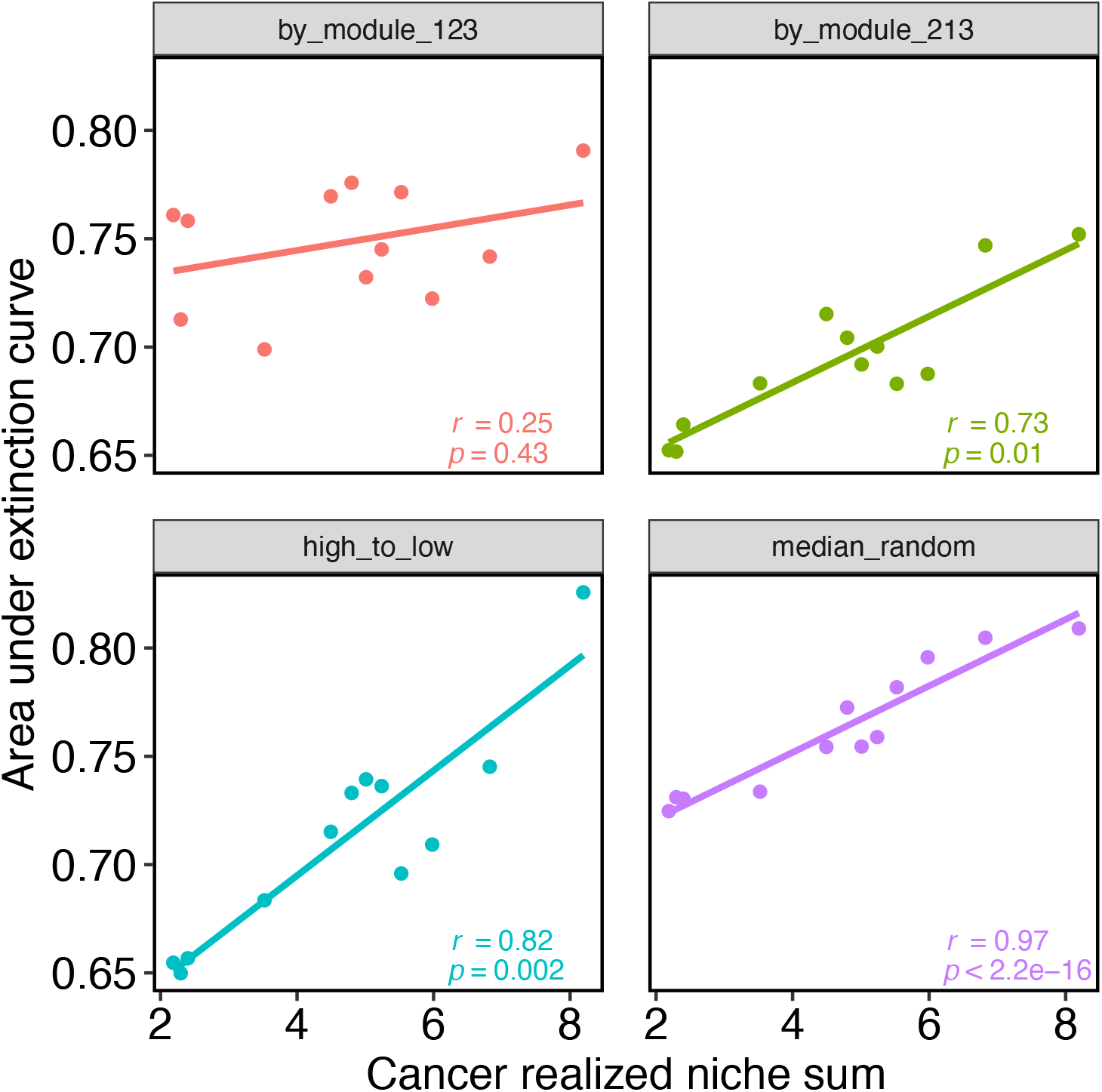
Correlation between robustness and realized niche. Each panel shows a scenario of node removal order. Each data point is a cancer type. A positive and significant correlation (values within the panels) indicates that cancer environments that enable chaperones to interact with a high proportion of the clients they can potentially interact with (high realized niche) will also be more robust to chaperone removal. Random removal serves as a control: we expect a strong positive correlation because when nodes are removed randomly, robustness is a function of network size and number of links, rather than of the nodes that are removed.

## Discussion

The mitochondria plays a crucial role in metabolic reprogramming of cancer cells [32]. The relationship between mitochondrial chaperones and the clients they interact with is key to cancer cell proliferation [33] but little is known about how CCI networks are structured and how this structure varies across cancer types. Portraying CCI networks as ecological multilayer networks of species interactions, we show for the first time that CCI network structure non-randomly depends on cancer type. In addition, we are able to identify the identity of chaperones divided into modules that provide redundancy for client support with consequences for network robustness. These results open the door for new hypotheses regarding the evolution of CCI networks and can inform caner-specific drug therapy drug development strategies.

We found a clear, non-random pattern of weighted nestedness in the number of clients chaperones interact with across cancers. Weighted nestedness requires variation and order in that variation in both chaperones and cancer types [23]. We did not detect an effect of chaperone or protein expression levels on chaperone’s realized niche. Hence, the mechanism underlying variation in realized niche (rows in Fig. 1) remains an open question for future research. A nested order can arise if core chaperones are crucial for supporting key proteins that must be sufficiently functional in all cancer types—as is the case of CLPP and HSPE1—while other chaperones are necessary only in particular cancer types to complement the function of clients supported by the core chaperones. Supporting this hypothesis is the result that chaperones with a high realized niche also tend to conserve their interaction partners across cancer types. A possible explanation for cancer-mediated variation (columns in Fig. 1) is the extent to which chaperones are a limiting factor. When expression does not have to be adjusted to meet the demand for supporting, there will be less co-expression correlations. Alternatively, it is possible that substrates of a chaperone are not exclusively dependent upon it except in a particular cancer type. From that variation, nestedness can then arise if there is a concomitant hierarchy in the importance of chaperones for client support.

Going beyond the number of clients interacted with, we found that chaperones are separated into distinct modules with regard to the clients they interact with. These modules imply matching, and most likely specialization of a group of chaperones on a group of clients. In community ecology, such pattern of limited niche overlap has been consistently found. For instance, plant-pollinator networks are often modular [34,35]. One mechanism underlying modularity is coevolutionary dynamics in which coevolutionary adaptations favor interactions with a set of species and exclude others [35]. Chaperone-client interactions are not mutualistic, excluding this hypothesis. However, selection operating at the cell-level can still favor cells in which protein that are being supported by more specialized chaperones results in more efficient interactions. This will result in modules of chaperones partitioned by their specialized clients. Another driver of modularity is trait matching. For instance, in pollination syndrome flower color can attract specific pollinators, generating modules [36]. In the world of proteins, modularity via trait matching can emerge if there is a match in the biophysical attributes of chaperones and their clients.

Selection is likely to favor the redundancy within each module because in case a chaperon becomes a limiting factor, another can take its place. The importance of redundancy to network robustness has been shown in ecological networks [37]. Another, non-mutually exclusive hypothesis is that chaperones within the same module have complementary functions because more than one is needed to complete the folding process [38,39]. Notably, SPG7 was the only chaperone in its group, and has the least number of interactions, making it a target for future research. The fact that the modules emerge across cancers indicates that the cancer environment has a small effect on selection pressures that underlie the evolution of trait matching. This result is strengthened by a previous study in which we used a single CCI network that ignored the cancer environment, and found a similar module composition [40].

Modularity is a structural feature directly related to network stability [20]. Here, robustness analysis revealed that in most cancer types the network collapses faster when chaperones are removed by their degree (high to low). This result is theoretically expected and has been shown before [18]. However, we also found that removing chaperones from module 2 resembled removal by degree, whereas removing from module 1 did not. Therefore, chaperones within module 2 are more important for network cohesiveness, making them ideal candidates for drug therapy targets. A striking finding is that module 2 contained a high proportion of co-chaperones. This suggests that co-chaperons are required for interacting with particular clients and therefore have the highest contribution to the network’s robustness. Moreover, when chaperones were removed in module 2 (but not module 1), there was a positive correlation between the extent to which a cancer environment supports protein interactions and robustness. Cancers in which chaperones interact with most of their clients are more robust to co-chaperone removal. Therefore, different cancer types may require the targeting of a different number of co-chaperones.

It has been argued that cancer research could benefit greatly from an ecological and evolutionary theories [10,11]. For instance, ecological theory can be used to order cancers by phenotypic axes in the “oncospace” [12]. This idea highlights the differences between cancers, supporting the cancer-dependent patterns we discover. Here, we apply for the first time theory and methodology from network ecology to study variation in protein interactions across cancer environments. Environmental variation in ecological interaction networks, and the link between structure and robustness are central themes in community ecology [9,17] and their application here provided novel insights into how chaperone interactions vary across cancer types, with consequences for robustness. This study can, in turn, inform ecological theory, where theory has been largely developed for communities of interacting organisms, not proteins. This is an important difference because in cancer, unlike in community ecology, the unit of selection are cells, or the hosts, rather than the interacting units. On the methodological side, nestedness has been extensively studied in the context of interactions [13,14,24], but not to detect patterns in the effect of environment on the realization of interactions.

Our study has several limitations. We inferred interactions from co-expression at the mRNA level. Nevertheless, genes may have evolved to be co-expressed for reasons different than a physical interaction or dependency, as is the case forthe P53 target gene induced by DNA damage [41]. Nevertheless, this remains a powerful method for estimating interactions [42]. We constructed binary networks in which an interactions exists or not. However, as has been shown in ecology, quantitative networks in which the strength of interaction is estimated may reveal different patterns and provide more insight into the importance of specific interactions [43]. A common limitation in ecological network analysis, including the one presented here, is that it is virtually impossible to quantify the importance of higher order interactions on each link [44]. Here, higher order interaction refers to the effect of a third client, chaperone or another cellular component on a chaperone-client interaction. For example, cooperation between chaperons such that a folding of a client depends on both. Robustness analysis is an in-silico approach that provides first insight into stability. Nevertheless, it does not capture the full spectrum of processes that operate in nature [45]. For instance, when a chaperone is removed, rewiring can occur such that another takes it place [46]. Moreover, we assumed that all proteins are equally necessary for tumor cells to operate but this assumption likely does not hold. In addition, the importance of proteins can also vary across cancers.

Our analysis provides only a starting point to understanding how CCI network structure affects its stability. Future computational studies that apply more realistic algorithms are needed to identify chaperons that can be used as targets for cancer therapy. In addition, incorporating the perspective of proteins would be necessary. These studies must be complemented with experiments that test chaperon removal. For example, one can use high throughput CRISPR and shRNA to identify chaperons and proteins which are essential for growth of cancer cell lines in culture [47,48]. The results we presented can serve as a starting point to guide these. From a theoretical perspective it would be beneficial to understand the evolutionary processes that drive cancer-depend variation in CCI networks.

## Materials and Methods

### Experimental Design

With the curation of the transcriptom data in the TCGA and making it available to the public, a door was opened to inquire about the way a cancer system as a whole performs, in the expression level, in the hope the insights will allow better treatment for the disease in the future.

In this study we used the gene expression level to assimilate a network for each cancer type using a correlations between expression levels. we used bootstraps to validate those correlations, and the resulting networks were used to be analysed.

### Data acquisition

Gene level transcriptome profiling (RNA-Seq) data (in the form of HTSeq - FPKM) was download from The Cancer Genome Atlas (TCGA) using the Genomic Data Commons Data Portal (https://portal.gdc.cancer.gov). Of these data we kept only the expression levels of mitochondrial proteins as listed according to MitoCarta 2.0 database.

### Co-expression analysis and network construction

Using the raw data of each cancer tissue, we calculated a Spearman correlation between the expression levels of 15 mitochondrial chaperone genes and 1,142 genes belonging to their potential protein substrates, using all available samples for that tissue. We created a chaperone-client interaction matrix (network). Chaperone-client pairs that were significantly correlated after Bonferroni correction for multiple testing received a value of 1, and non-significant correlations received a 0 (no interaction).

Cancer types varied in their sample size (Table S1), which can create biases in the statistical power for detecting significant correlations. To ensure a fair comparison between all cancer types we performed a bootstrap analysis to match the sample size of each cancer type to that with the least number of samples (Kidney Renal Papillary Cell Carcinoma, *n* = 288). For each cancer type we drew 288 transcriptome samples at random without replacement (1,000 attempts), and reconstructed the chaperone-client network using Spearman correlations (as described above). A correlation that was classified as significant and positive in at least 95% of the bootstrapping attempts was considered as an interaction. Results of this analysis are in Table S1.

### Nestedness analysis

We calculated weighted nestedness as the largest eigenvalue of a matrix, *ρ* [24]. This method is particularly suitable for comparing a matrix to its shuffled counterparts (see below for details on shuffling). Specifically, given a set of weighted interactions, the matrix with the highest weighted nestedness is that in which the distribution of matrix cell values produces the largest *ρ* [24].

### Modularity analysis

We conducted a modularity analysis to detect groups (modules) of chaperones and clients that interact densely with each other. For this analysis we used the 12 cancer types together, effectively creating a multilayer network in which each layer was a chaperone-client interaction network of a given cancer type. We detected the optimal partitioning to modules using Infomap [28,49]. Briefly, Infomap detects an optimal network partition based on the movement of a random walker on the multilayer network (see [28,49,50] for details). For any given partition of the network, the random walker moves across nodes in proportion to the weight of the edges. Hence, it will tend to stay longer in dense areas. These areas can be defined as ‘modules’. The movement dynamics can be converted to an information-theoretic currency using the objective function *L* called the map equation. The optimal network partition corresponds to that with the minimum value of *L* [49]. In multilayer networks, the random walk also moves from a node in one layer (e.g. HSPD1 in BRCA) to its counterpart in another layer (e.g. HSPD1 in KIRP) with a given rate called the relax rate. This connects the layers together, allowing Infomap to find modules of chaperones and clients that are connected across cancer types. See details on Infomap and how it is applied to ecological multilayer netowrks in [28].

### Network shuffling

We wanted to ensure that the network properties we find for empirical networks (e.g., nestedness, similarity in interactions, Jaccard indices) are not a result of random processes but rather are biologically significant. Hence, we compared the results of empirical networks to those obtained from analyses of 1,000 counterpart shuffled networks. This is a common procedure in the study of ecological networks [51–53]. We shuffled the interaction values of each CCI matrix using the ‘curveball’ algorithm [54], which constrains the number of client associations (for chaperones) and chaperone associations (for proteins) to that observed in the empirical networks. This algorithm is highly conservative making it difficult to detect statistically significant results; or in other words, increases the likelihood of type II error [55]. This increases our confidence that the statistically significant results we find are true biological processes.

Using the shuffled networks, we determined statistical significance of weighted nestedness using a one-tailed test as is common in ecological networks [24,52]. We compared the empirical *ρ* to its shuffled counterparts using the following formula:

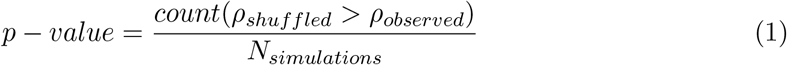

To calculate the statistical significance of measures at the node level (i.e., the Jaccard index calculations), we used z-scores as follows [26].

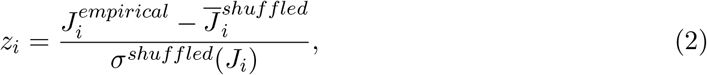

where 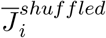 and *σ^shuffled^*(*J_i_*) are the mean and standard deviation of the Jaccard index obtained from the shuffled networks. Hence, a positive (negative) z-score suggests that similarity is higher (lower) than expected from random CCIs. Significance of each empirical value was determined at the 0.05 level: a z-score > 1.96 or < −1.96 indicate that the index is greater or lower than the random expectation, respectively.

## Code and data availability

All analyses were done in R (version 4.1) within the Linux environment. Network analysis was done using the packages ‘bipartite’ (2.1636), ’vegan’ (2.5-7) and infomapecology. All the code and the final binary matrices are available in https://github.com/Ecological-Complexity-Lab/cancer_neworks.

## Acknowledgements

The data curation in this project was done with the help of Dr. Liron Levin.

## Data and materials availability

All data, code, and materials used in the analyses will be available in GitHub upon acceptance.

## Supplementary Materials

**Table S1:** Summary of results for normalizing the significant chaperone-client correlations by number of samples in each cancer. For each cancer tissue specified are: the number of samples of each cancer cohort, the number of co-expression correlations found to be significant using spearman correlations, the number of correlations found to be valid after bootstrapping (see Methods), and the percent of correlation kept after bootstrapping out of the number of the significant correlations using spearman alone.

| Abbreviation | Cancer | Samples | All Correlations | N valid | % valid |
| --- | --- | --- | --- | --- | --- |
| KIRP | Kidney renal papillary cell carcinoma | 288 | 5108 | 5108 | 100% |
| LIHC | Liver hepatocellular carcinoma | 371 | 3693 | 2148 | 58.16% |
| STAD | Stomach adenocarcinoma | 375 | 5015 | 3083 | 61.48% |
| COAD | Colon adenocarcinoma | 478 | 5665 | 2840 | 50.13% |
| PRAD | Prostate adenocarcinoma | 498 | 6236 | 3661 | 58.71% |
| HNSC | Head and Neck squamous cell carcinoma | 500 | 7466 | 3612 | 48.38% |
| LUSC | Lung squamous cell carcinoma | 502 | 4641 | 1463 | 31.52% |
| THCA | Thyroid carcinoma | 502 | 6945 | 4305 | 61.99% |
| LUAD | Lung adenocarcinoma | 533 | 4577 | 1522 | 33.25% |
| KIRC | Kidney renal clear cell carcinoma | 538 | 6242 | 3089 | 49.49% |
| UCEC | Uterine Corpus Endometrial Carcinoma | 551 | 6461 | 3201 | 49.54% |
| BRCA | Breast invasive carcinoma | 1102 | 6358 | 1411 | 22.19% |

**Table S2:**
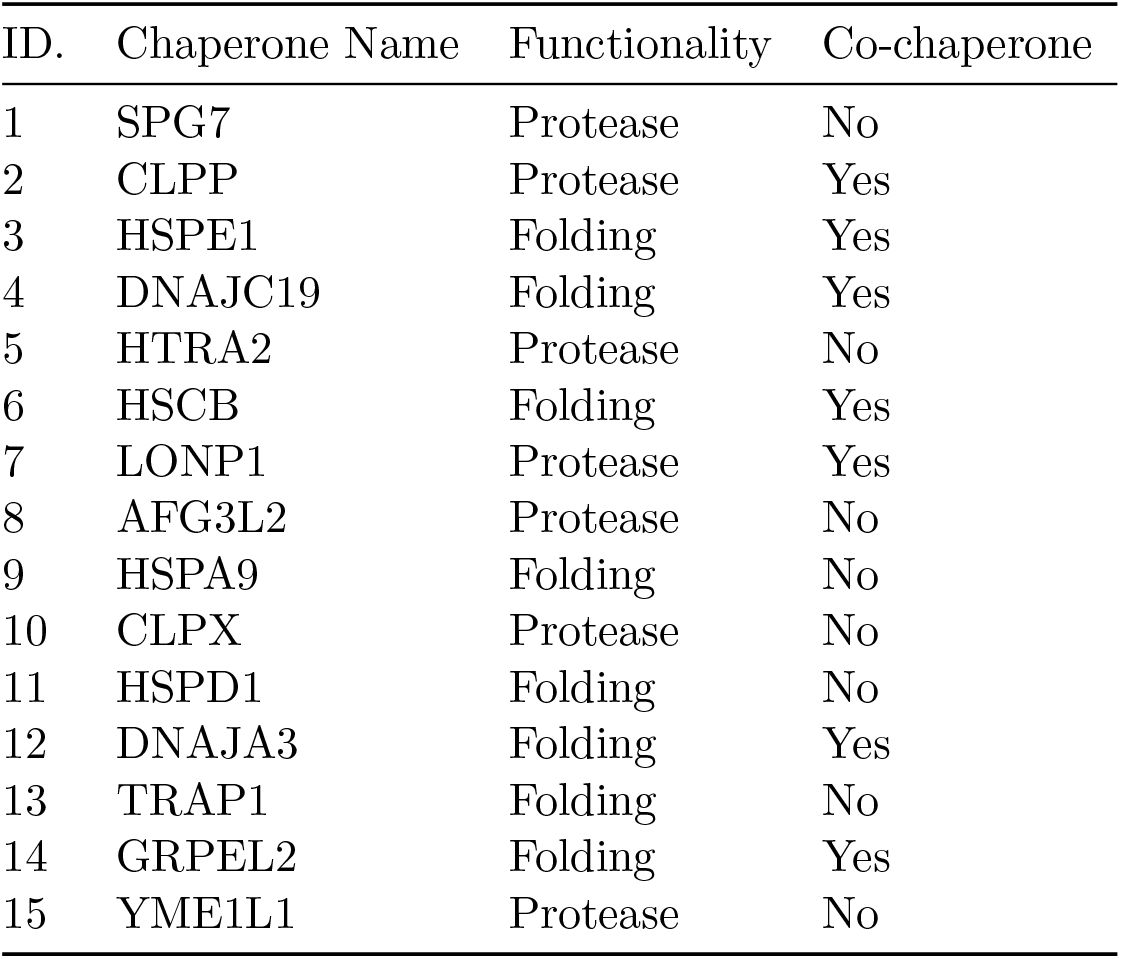
List of the chaperones. For each chaperone we specified: an ID (used in fig. 4B); its corresponding gene name; main biological function and whether it is a co-chaperone.

| ID. | Chaperone Name | Functionality | Co-chaperone |
| --- | --- | --- | --- |
| 1 | SPG7 | Protease | No |
| 2 | CLPP | Protease | Yes |
| 3 | HSPE1 | Folding | Yes |
| 4 | DNAJC19 | Folding | Yes |
| 5 | HTRA2 | Protease | No |
| 6 | HSCB | Folding | Yes |
| 7 | LONP1 | Protease | Yes |
| 8 | AFG3L2 | Protease | No |
| 9 | HSPA9 | Folding | No |
| 10 | CLPX | Protease | No |
| 11 | HSPD1 | Folding | No |
| 12 | DNAJA3 | Folding | Yes |
| 13 | TRAP1 | Folding | No |
| 14 | GRPEL2 | Folding | Yes |
| 15 | YME1L1 | Protease | No |

**Fig. S1:**
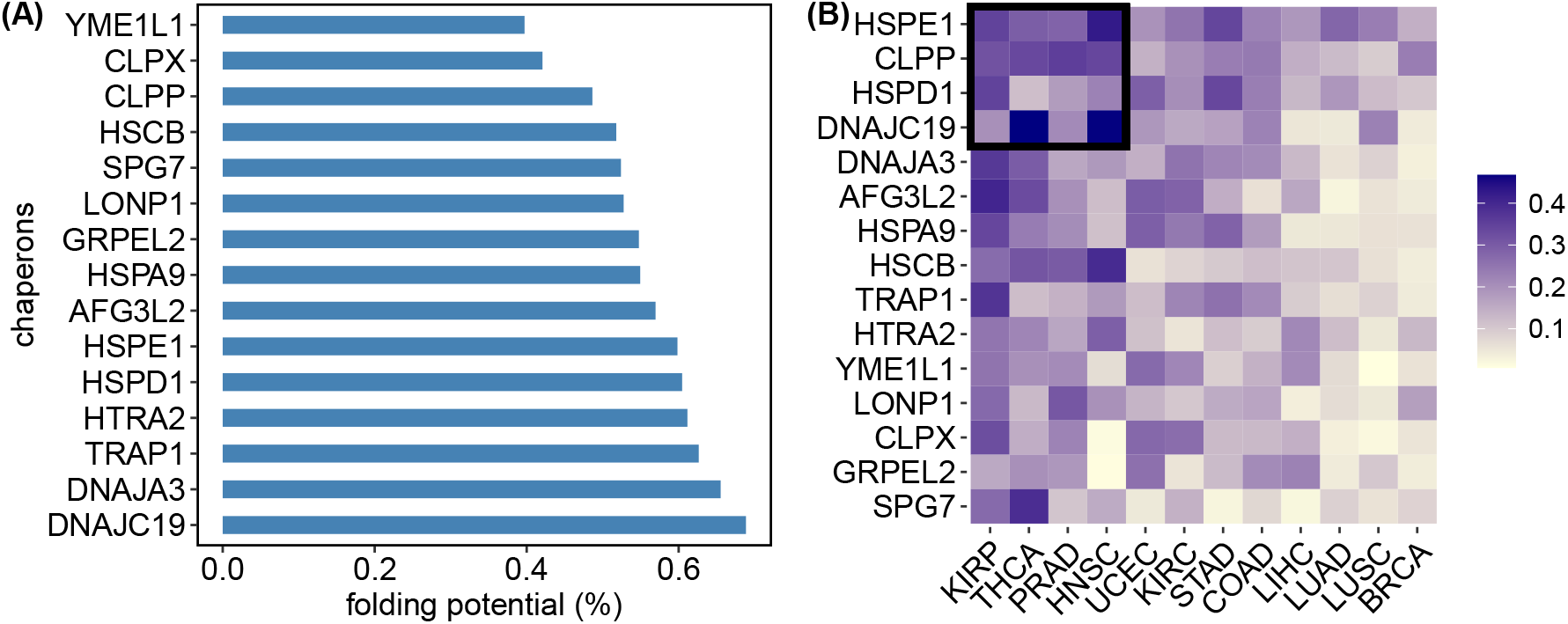
chaperone specialization. **(A)** Specialization *S_c_* is a proportion calculated as the total number of clients that a chaperone interacts with across cancers, out of all the 1,142 mitochondrial proteins.**(B)** Each square in the heatmap depicts a chaperone’s cancer-specific specialization, 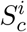, defined as the number of clients that a chaperone (rows) interacts with in a given cancer environment (columns), 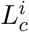, out of all the proteins (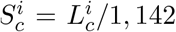; see Results). The non-uniform colors in each row indicate that each chaperone interacts with a different number of clients in across cancers. Furthermore, the non-uniform colors in each column indicate that cancer types also vary in the extent to which they enable chaperones to interact with clients. Put together, these two observations create a weighted-nested pattern whereby the more specialized chaperones interact with less clients and are a subset of the more generalist ones; on the other hand, cancer environments that enable chaperones to interact with less clients are subsets of those that allow for higher levels of generalism. Weighted-nestedness was statistically significantly non-random when compared to 1,00 counterpart networks assembled from networks in which chaperone-clients interactions were shuffled (Methods; Fig. S2A). Therefore, there is a non-randomly structured way in which cancer environments mediate the interactions of chaperones. Rows and columns are arranged by their marginal sums.

**Fig. S2:**
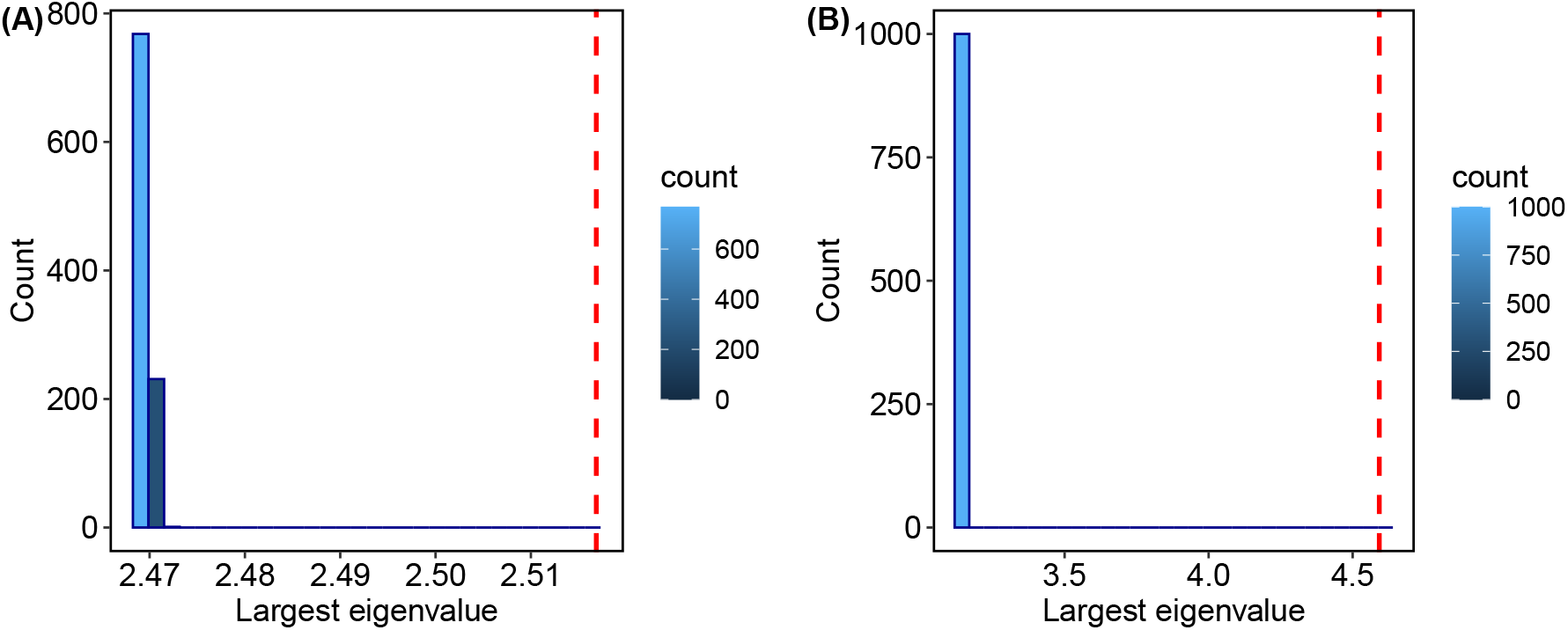
Significance of weighted nestedness. Nestedness was calculated as the largest eigenvalue of each matrix in Fig. 1 in the main text. This value was compared to a distribution of 1,000 values calculated for shuffled networks. The observed value (dashed vertical line was larger than all values obtained for shuffled networks for specialization **(A)** and realized niche **(B)**.

**Fig. S3:**
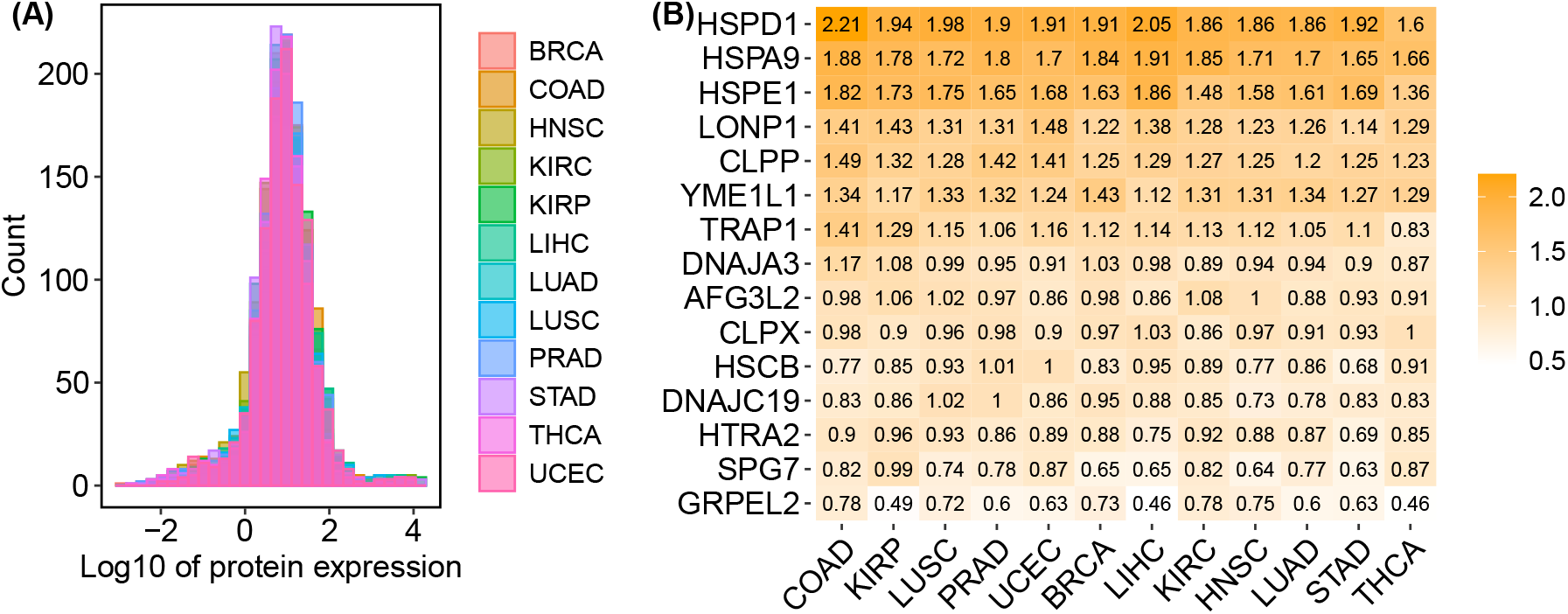
Gene expression levels are similar across cancers but vary across chaperones. **(A)** The median expression value of each mitochondrial protein was calculated and log-transformed for comparable expression scales. Each color depicts the distribution of these medians in a given cancer. **(B)** The median value for each chaperone’s expression level in each cancer type was calculated, and transformed by log10 to compare expression scale. Each value is presented in the corresponding cell in the heatmap. Rows and columns are arranged by their marginal sums.

**Fig. S4:**
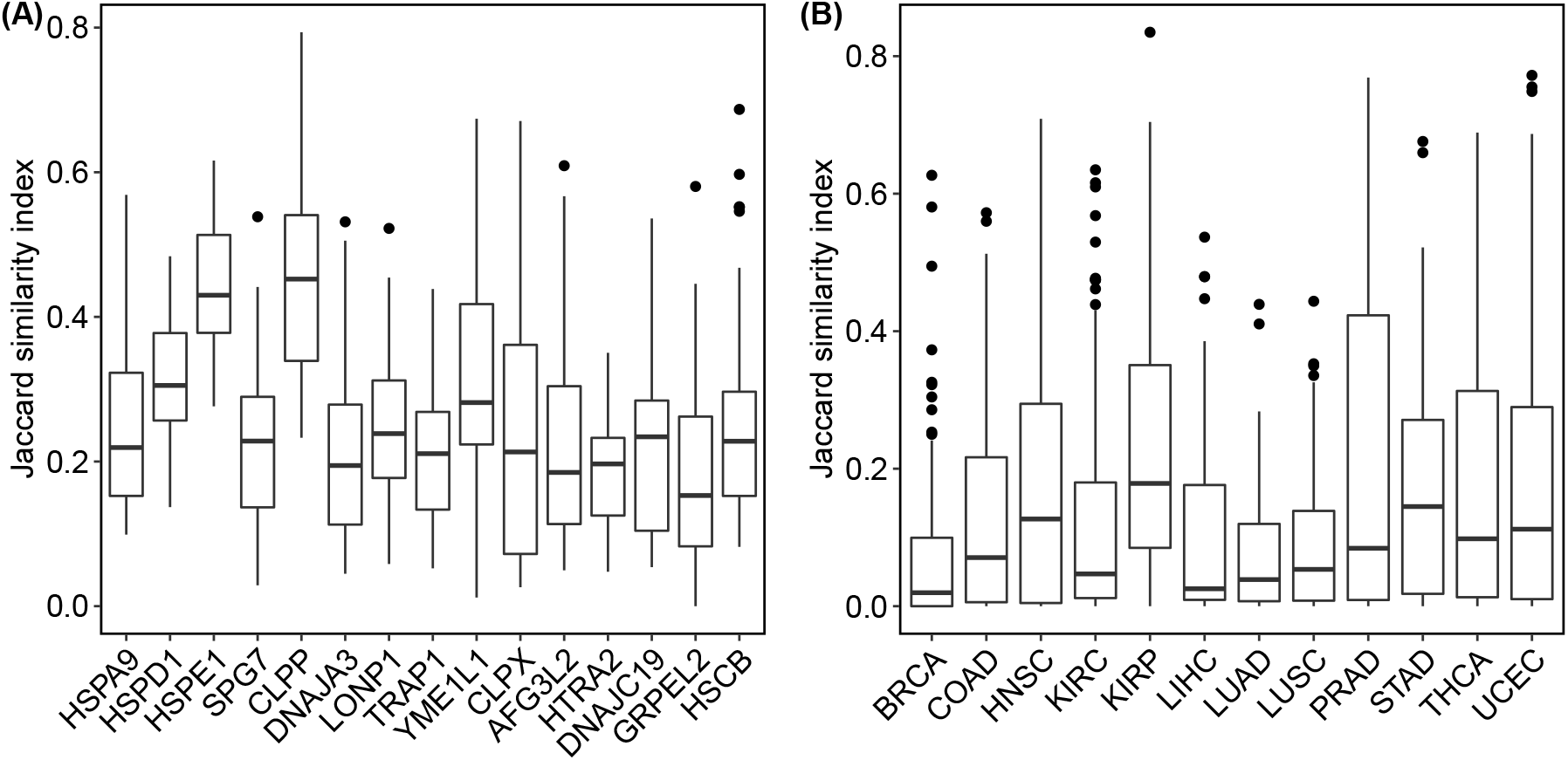
Jaccard similarity index distributions. **(A)** Distribution of Jaccard similarity for the identity of clients for each chaperone *c* between all pairs of cancer types *i* and 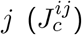. **(B)** Distribution of Jaccard similarity for the identity of clients for every pair of chaperones *x* and *y*, within each cancer type 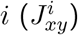.

**Table S3:** Spearman correlation values for the correlations of realized niche with chaperone expression for each chaperone. The Bonferroni-corrected significant p-value is 0.003

| chaperone | p-value | R-value |
| --- | --- | --- |
| AFG3L2 | 0.931 | 0.028 |
| CLPP | 0.753 | 0.102 |
| CLPX | 0.404 | -0.266 |
| DNAJA3 | 0.404 | -0.266 |
| DNAJC19 | 0.746 | -0.105 |
| GRPEL2 | 0.145 | -0.448 |
| HSCB | 0.829 | -0.0699 |
| HSPA9 | 0.297 | -0.329 |
| HSPD1 | 0.527 | 0.203 |
| HSPE1 | 0.255 | -0.357 |
| HTRA2 | 0.276 | -0.343 |
| LONP1 | 0.966 | 0.014 |
| SPG7 | 0.131 | 0.462 |
| TRAP1 | 0.379 | 0.280 |
| YME1L1 | 0.00824 | -0.720 |

